# Truncated, uncapped mRNA 5’ ends targeted by cytoplasmic recapping cluster at CAGE tags and some transcripts are alternatively spliced

**DOI:** 10.1101/471813

**Authors:** Mikaela R. Berger, Rolando Alvarado, Daniel L. Kiss

## Abstract

Until cytoplasmic recapping was discovered, decapping was thought to irreversibly destine an mRNA to degradation. Contradicting this idea, we readily observe mRNAs targeted by cytoplasmic capping in uncapped, yet stable forms. 5’ RACE shows that nearly all uncapped ends correspond to CAGE tags and that the recapping of ZNF207 mRNA may be restricted to a single splice isoform. A modified RACE approach detected uncapped 5’ RNA ends mapping to 46 mRNAs in dominant negative cytoplasmic capping enzyme expressing and normal cells. 11 of 46 cloned mRNAs also contained splice isoform-limiting sequences. Collectively, these data reinforce earlier work and suggest that alternative splicing may play a role in targeting transcripts for– and/or determining the position of– cytoplasmic capping.

## Introduction

The N7-methylguanosine (m^7^G) cap is a critical modification added to the 5’ ends of mRNAs during transcription. Nuclear capping occurs in three steps. First, the triphosphatase domain of RNA guanylyltransferase and 5’-phosphatase (RNGTT, capping enzyme hereafter) removes the gamma phosphate from the first transcribed nucleotide [1]. Capping enzyme then transfers an inverted guanosine residue onto the nascent RNA using its guanylyl transferase domain [1]. Finally, the cap is methylated by RNA guanine-7 methyltransferase (RNMT) [2]. The m^7^G cap is vital for proper mRNA splicing, processing, packaging, export, and translation [3, 4]. The m^7^G cap is a focal point of different RNA surveillance pathways and helps protect the mRNA from cellular 5’ to 3’ exonucleases [4]. Although evidence for uncapped mRNA species with mono- and di-phosphate 5’ ends dates back to the 1970’s [5, 6], until recently, decapping was generally thought to irreversibly commit an mRNA to degradation [7, 8].

Certain nonsense-mediated decay (NMD) decay intermediates generated from premature termination codon containing β-globin mRNAs were found to violate this rule [9, 10]. These 5’-truncated mRNA decay intermediates were stable, accumulated in a capped form [9], and required a functional NMD pathway to form [11]. Since NMD occurs exclusively in the cytoplasm, these data alluded to a cytoplasmic RNA recapping mechanism [3, 9]. These capped β-globin decay intermediates remained an unexplained oddity for many years; however, the advent of reliable transcriptome-wide methods to assay different RNA populations uncovered the existence of different uncapped RNA species in plants [12, 13] and in mammals [14]. Finally, capped analysis of gene expression (CAGE), a technique designed to identify transcription start sites by tagging the position of m^7^G caps on mRNAs, showed that nearly 25% of mammalian m^7^G caps were not at known transcription start sites [15]. Rather, they were located within the body of a transcript and the authors hypothesized mRNAs could be recapped after cleavage or truncation [15].

Cytoplasmic capping was first described around this time [16]. That work identified an RNA capping activity in cytoplasmic extracts and observed that blocking that activity by overexpressing a dominant negative cytoplasmic capping enzyme (K294A) inhibited the cell’s ability to recover from stress [16]. The cytoplasmic capping complex forms when Nck1 interacts with a monophosphate kinase and capping enzyme to coordinate the first two steps of cytoplasmic capping (Figure 1A) [17]. The newly added cap is then methylated by RNMT [18]. Caps added in the cytoplasm appear to be indistinguishable from nuclear-added caps. Roughly 2000 mRNAs are targeted by cytoplasmic capping and that these mRNAs are less frequently associated with translating polysomes when K294A is expressed [19]. Recent work has also shown that uncapped cytoplasmic capping targets retain translationally functional poly(A) tails [20] and that 5’ RACE can detect uncapped ends from 5’ truncated mRNAs in the vicinity of CAGE tags in cells expressing K294A [21]. Collectively, these findings implied that novel 5’ ends from truncated mRNAs should be observable among the non-translating mRNAs in cells where cytoplasmic capping was unimpeded in addition to those where cytoplasmic capping had been blocked.

**Figure 1.**
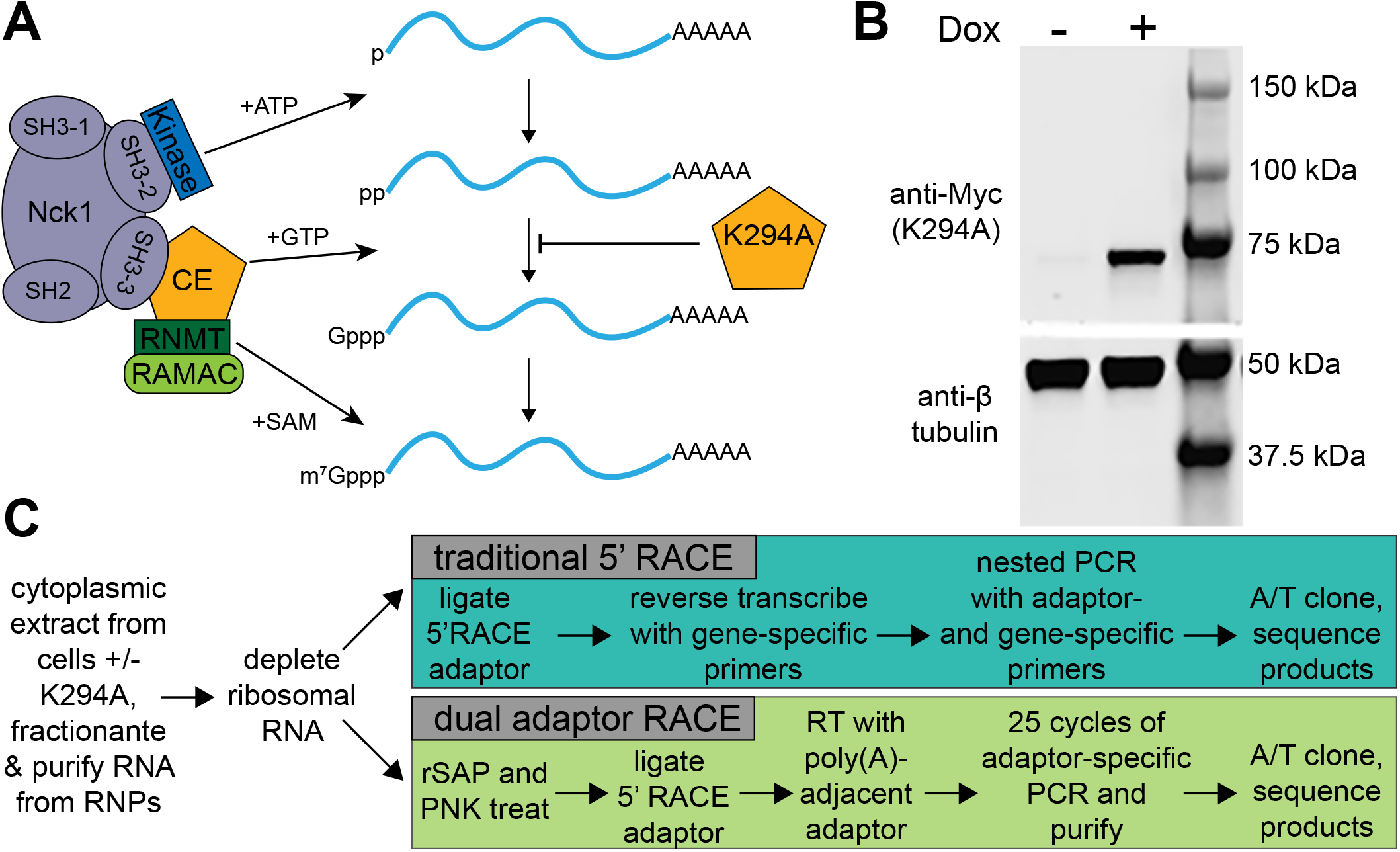
Utilizing a dominant negative capping enzyme (K294A) to study cytoplasmic capping. **(A)** A schematic demonstrating the organization of, and activities within, the cytoplasmic capping complex. **(B)** Cytoplasmic extracts were harvested (see methods) from uninduced (−Dox) and K294A-expressing U20S cells (+Dox) and assayed via western blotting. A western blot probed with anti-Myc antibody shows inducible expression of K294A from a representative pair of samples. The bottom portion of the same blot was probed with an anti-tubulin antibody and serves as a loading control. **(C)** A flow chart showing the workflow for the traditional 5’ RACE (blue) and dual adaptors RACE (green) arms of this two-armed study.

In this report, we readily detected uncapped and 5’ truncated mRNAs in cytoplasmic extracts from uninduced cells, where cytoplasmic capping was normal, in addition to those expressing K294A. As before, nearly all of the uncapped ends mapped to CAGE tags. While most of the putative recapping sites yielded sequences that could not discriminate between different splice isoforms, surprisingly, one cluster of 5’ ends was specific to a single isoform of ZNF207 mRNA. To better evaluate this intriguing possibility, we used a dual adaptor RACE approach to identify additional uncapped mRNA ends. Upon closer examination, almost one in four of the cloned mRNA ends contained sequences specific to a limited number of splice isoforms. These findings offer the first hints that sequence cues may help determine the identity of– and/or the position of– cytoplasmic mRNA recapping.

## Materials and Methods

### Cell culture and preparation of cytoplasmic extracts

K294A cells generated in [16] were cultured in McCoy’s medium (Gibco) supplemented with 10% tetracycline-tested fetal bovine serum. 150 mm tissue culture dishes were seeded with 3 × 10^6^ log-phase cells. 24h later, expression of K294A was induced by adding doxycycline (1 μg/ml). 24h later, cells were rinsed twice with phosphate buffered saline, scraped from plates with cell lifters, and pelleted by centrifugation at 2500 ×g for 5 min. Cytoplasmic extracts were prepared by lysing pellets in 5 volumes of ice cold cytoplasmic lysis buffer (50 mM Tris-HCl pH 7.5, 10 mM KCl, 10 mM MgCl2, 150 mM NaCl, 0.2 % NP-40, 2 mM DTT, 0.5 mM PMSF, 1 mM sodium orthovanadate, supplemented with 5 μl/ml RNAseOUT (Life Technologies), 25 μl/ml protease inhibitor cocktail (Sigma), 10 μl/ml each phosphatase inhibitor cocktails 2 and 3 (Sigma)) followed by incubation on ice for 10m with gentle agitation every 2m. Nuclei were removed by centrifuging at 16100 ×g for 10m at 4°C and supernatants were used for western blotting and cytoplasmic RNA isolation.

### Western blotting

Western blots were performed as in [20]. Cytoplasmic extracts were heated to 95°C in 1x Laemmli sample buffer for 5 minutes and then separated on 10% Mini-PROTEAN^®^ TGX^TM^ gels (Bio-Rad) and transferred onto Immobilon-FL PVDF membranes (Millipore). Membranes were blocked with 1% milk in Tris-buffered saline containing 0.05% Tween-20 (TBS-T), and incubated with rabbit anti-MYC or anti-Tubulin polyclonal antibodies (1:2500 dilution) in 0.1% nonfat milk overnight at 4°C. Blots were washed with TBS-T, incubated with LI-COR IR-680 anti-rabbit secondary antibody (1:10000 dilution) for 2 hours, washed with TBS-T and visualized using Odyssey V.3.0.30 software (LI-COR).

### Oligonucleotides used in this study

All oligonucleotides used in this study are shown in Table S1.

### Isolation and preparation of cytoplasmic RNA

RNA was harvested from cytoplasmic extracts using Direct-Zol kits (Zymo) according to the manufacturer’s reaction clean-up protocol (including DNase I digestion). 5 μg of cytoplasmic RNA was treated with Ribo-Zero Gold (Illumina), and all RACE experiments used the rRNA-depleted equivalent of 1 μg of cytoplasmic RNA.

### 5’ rapid amplification of cDNA ends (RACE)

Standard 5’ RACE was performed exactly as described in [21]. 25% of the ligation reaction was reverse transcribed (Superscript III, Life Technologies) using the manufacturer’s gene specific primer protocol (Table S1). After RNase H treatment, cDNAs were amplified using nested PCR with the indicated (Table S1) primers. Primary nested PCR products were purified using DNA Clean and Concentrator-5 columns (Zymo). Secondary PCRs were performed using CAGE site targeting primers (Table S1) as in [21], column purified, cloned into the pGEM-T-easy vector system (Promega), transformed, plated onto plates containing ampicillin, Isopropyl β-D-1-thiogalactopyranoside (IPTG), and 5-bromo-4-chloro-3-indolyl-β-D-galactopyranoside (X-Gal), and incubated at 37°C overnight. White colonies were assayed by colony PCR using primers complimentary to T7 and SP6 promoter sequences (Table S1). Products were separated using 2% agarose gels and ~300+ bp inserts were Sanger sequenced at the OSU Genomics Shared Resource. Sequences lacking RACE adaptors were discarded, and adaptor-containing sequences were identified using NCBI’s Basic Local Alignment Search Tool (BLAST) and splice variants were assigned using NCBI databases.

### CAGE databases

CAGE tags mapping to the RefSeq annotated transcription start site and cloned 5’ RACE ends of ITGB1, SARS, and ZNF207 mRNAs were mined from the following predefined track: [FANTOM CAGE TSS] All FANTOM5 CAGE libraries (n=1897 pooled) the week of Oct 15, 2018 using the ZENBU genome browser [22]. Only CAGE tags mapping to the sense strand were retained.

### Reverse transcription, qPCR and standard RT-PCR

1 μg of cytoplasmic RNA harvested from induced and uninduced K294A cells was spiked with 0.1 ng of luciferase control mRNA (Promega) and reverse transcribed with Superscript III (Life Technologies) using the manufacturer’s random priming protocol. qPCR reactions used SsoAdvanced Universal SYBR Green Supermix as instructed by the supplier (Bio-Rad) and data were analyzed via CFX Maestro software (Bio-Rad). Traditional RT-PCRs used gene specific primers with 2X Hot Start Master Mix (Apex) and were separated on agarose gels and visualized with a Gel Doc (Bio-Rad). All amplicons for qPCR and RT-PCR were validated by purifying bands from agarose gels and Sanger sequencing.

### Dephosphorylation and phosphorylation of RNA ends

5 μg of mRNA harvested from the mRNP fractions of polysome gradients harvested during [20] was Ribo-Zero Gold-depleted and 50% of the resulting RNA was heated to 65°C for 5m and flash cooled on ice. Samples were treated with recombinant shrimp alkaline phosphatase (rSAP) (NEB). To minimize precipitation steps, rSAP treatment used 1x polynucleotide kinase (PNK) buffer, where rSAP activity is reduced, therefore, the reaction time was increased to 3h at 37°C. rSAP was inactivated by adding EDTA and heating to 65°C for 5m. Mixtures supplemented with additional MgCl2, 10x PNK buffer, ATP, and PNK. Samples were incubated for 1h at 37°C and reactions were stopped with EDTA and heat denaturation. Kinased RNAs were ethanol precipitated and resuspended in water.

### Dual adaptor RACE

The dual adaptor RACE protocol utilizes rSAP and PNK treated RNA (see above) and was performed as standard 5’ RACE until the ligation step was completed. 25% of the ligation reaction was primed with the poly(A) adjacent adaptor (Table S1) and reverse transcribed with Superscript III (Life Technologies) according to the manufacturer’s instructions for oligo dT priming. cDNA was RNase H digested and PCR amplified (25 cycles) using primers targeting the 5’ and 3’ adaptor sequences (Table S1). PCR products were purified, ligated, transformed, plated, and colonies were screened and sequenced as above.

## Results and Discussion

### Most uncapped 5’ ends for ITGB1, SARS, and ZNF207 mRNAs support earlier observations and map to CAGE tags

Previous works have shown that uncapped forms of mRNAs targeted by cytoplasmic recapping were enriched in non-translating RNA pools [19], retain translationally functional poly(A) tails [20], and had uncapped 5’ mRNA ends in the vicinity of CAGE tags [21] when cytoplasmic capping was blocked. A large population of uncapped yet stable mRNAs, called natively uncapped mRNAs, were also found when cytoplasmic capping was normal [19]. We reasoned that these novel 5’ mRNA ends should be observable among non-translating mRNAs in cells where cytoplasmic capping was unimpeded as well as those expressing K294A (Figure 1B). Three mRNA targets, ITGB1, SARS, and ZNF207, were revisited with more detailed 5’ RACE experiments to evaluate this possibility [20].

In total, 98 positive (15 for ITGB1, 35 for SARS, and 48 for ZNF207) clones were obtained, 51 from uninduced and 47 from K294A-expressing cells, which mapped to 28 different truncated 5’ ends (Figure 2, Table S2). The recovery of uncapped mRNA ends in uninduced cells, where cytoplasmic capping is normal, and K294A-expressing cells confirms earlier published results and shows that a pool of uncapped mRNAs are present in normal cells [19]. When examined closely, 41 of the cloned 5’ RACE ends (Figure 2, Table S2) exactly matched the previously reported 5’ RACE ends from K294A-expressing cells [21]. Although not exact matches, 21 additional clones map within 7nt of previously reported uncapped 5’ ends, meaning that nearly two-thirds of these 5’ RACE clones agree with previously published data [21]. Of the assayed mRNAs, ITGB1 showed the largest difference when detecting downstream uncapped 5’ ends in K294A-expressing or uninduced cells (1 vs 14 clones) (Figure 2, Table S2). In contrast, downstream 5’ ends for SARS and ZNF207 mRNAs were more readily detectable in cells where cytoplasmic capping was blocked (20 vs 15 clones and 26 vs 22 clones respectively, Table S2) than in uninduced cells. As expected, these results show that different uncapped mRNA species behave differently *in vivo* although these data are insufficient to suggest a possible regulatory mechanism to account for the observed differences.

**Figure 2.**
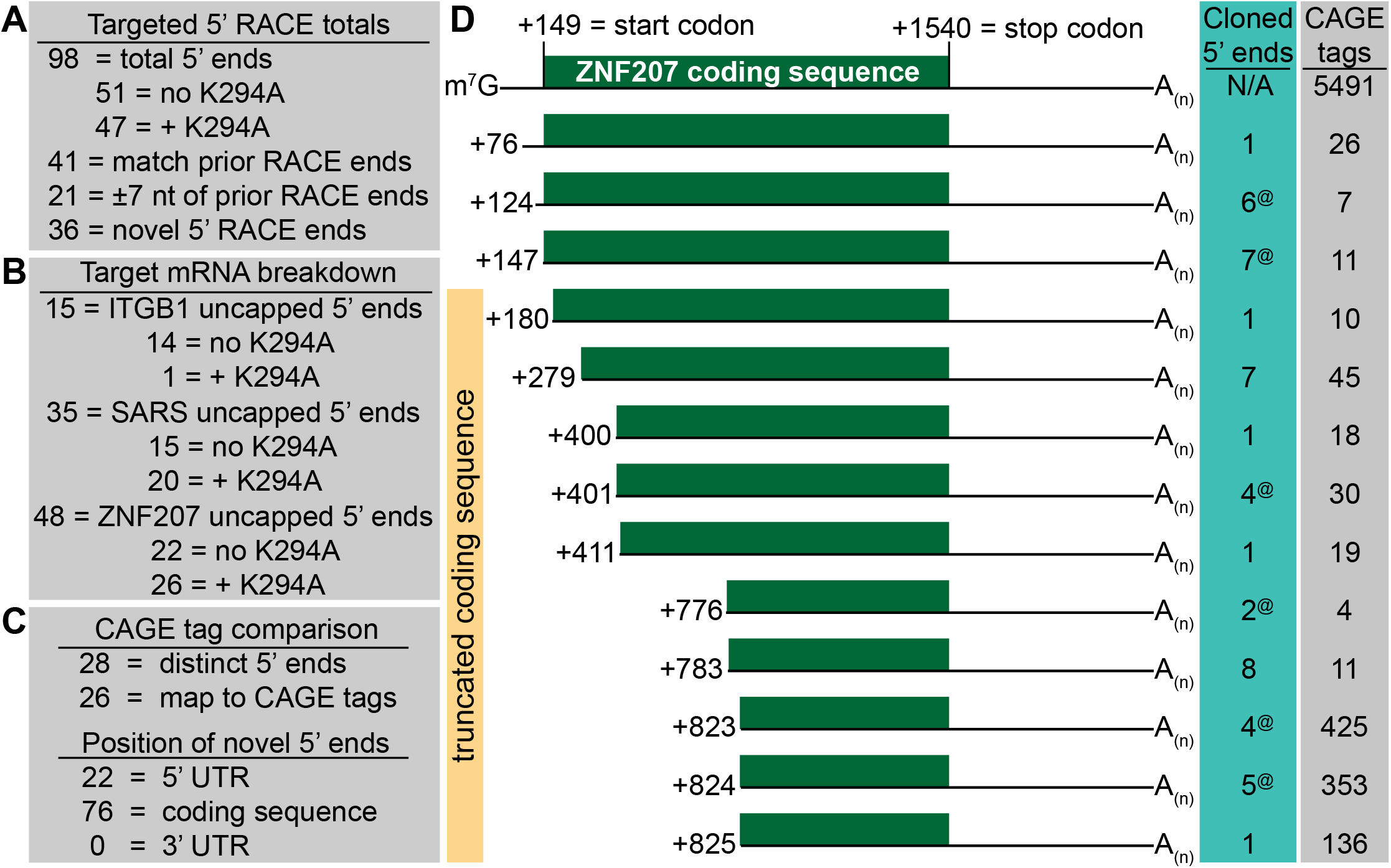
The uncapped 5’-ends of truncated cytoplasmic capping-targeted mRNAs correspond to CAGE tags. Three mRNAs known to be cytoplasmic capping targets were assayed using 5’ RACE. **(A)** The total number of novel 5’ ends detected, their overlap with, or proximity to, previously published 5’ RACE data are indicated. **(B)** The number of 5’ truncated mRNA ends mapping to each assayed cytoplasmic capping target are shown. Whether they were detected in the presence or absence of K294A is also indicated. **(C)** The overlap between the 5’ ends identified in this work and previously published CAGE tags plus the positions of the 5’ RACE ends in regards to the coding sequence of their mRNAs are indicated. **(D)** Schematics showing the positions of the truncated 5’ ends cloned by 5’ RACE relative to the full-length ZNF207 mRNA. The green box in each schematic represents the coding sequence of the mRNA. The full-length mRNA (top) is indicated by m^7^G, and the nucleotide positions (for isoform 2) of the novel, 5’-truncated ends are shown to the left of each schematic. The number of times each 5’ end was cloned (blue box) and the number of CAGE tags mapping to that base (grey box) are indicated to the right of each schematic. ^@^ denotes an exact 5’ RACE match to uncapped ends published in [21].

The previous work also mined 18 CAGE libraries to determine if these uncapped 5’ ends correlated with published CAGE tags. This analysis was revisited by using the Zenbu interface for Phantom5 to mine (see methods) nearly 1900 CAGE libraries (Figures 2, S1) for CAGE tags mapping to these three mRNAs [22]. In total, 26 of the 28 downstream, uncapped 5’ ends for ZNF207, SARS and ITGB1 mRNAs detected by these targeted 5’ RACE experiments map to documented CAGE tags (Figures 2, S1). This means that nearly all of these 5’ ends correspond to cap positions distinct from the annotated transcription start site and several are inside the coding sequences of their respective mRNAs (Figures 2, S1). While downstream transcription initiation events can’t be ruled out as the source of the observed CAGE tags, such events are unlikely to be the direct source of the novel 5’ ends observed here. This is because mRNAs are capped co-transcriptionally, meaning that they would need to be decapped prior to being identified in our 5’ RACE protocol.

### An alternatively spliced isoform of ZNF207 mRNA is preferentially recovered by 5’ RACE

A closer examination of the 5’ RACE products showed that seventeen of twenty 5’ RACE clones near the middle of ZNF207 mRNA included exon 8/10 junction sequences (Table S2). The three predominant isoforms of ZNF207 mRNA have been shown to differ by their inclusion of exons 6 and 9 (Figure 3A) with exon 8/10 junctions exclusively present in isoform 2 [23]. Importantly, the three outliers were too short and lacked that isoform-defining junction (Figure 3A, Table S2). One possible explanation of this result was that only isoform 2 of ZNF207 mRNA was expressed in U20S cells. Traditional RT-PCR experiments show that both isoforms 2 and 3 of ZNF207 mRNA are expressed while isoform 1 is absent in U20S cells (Figure 3B). These RT-PCR experiments also show that isoform 2 is roughly twice as common as isoform 3 and that the total levels of ZNF207 mRNA decreased slightly in response to K294A expression (Figure 3B). The latter result was validated with qPCR experiments which also show a ~20% reduction in total ZNF207 mRNA levels upon K294A treatment. Further, both expressed isoforms of ZNF207 are similarly decreased upon K294A induction (Figure 3C).

**Figure 3.**
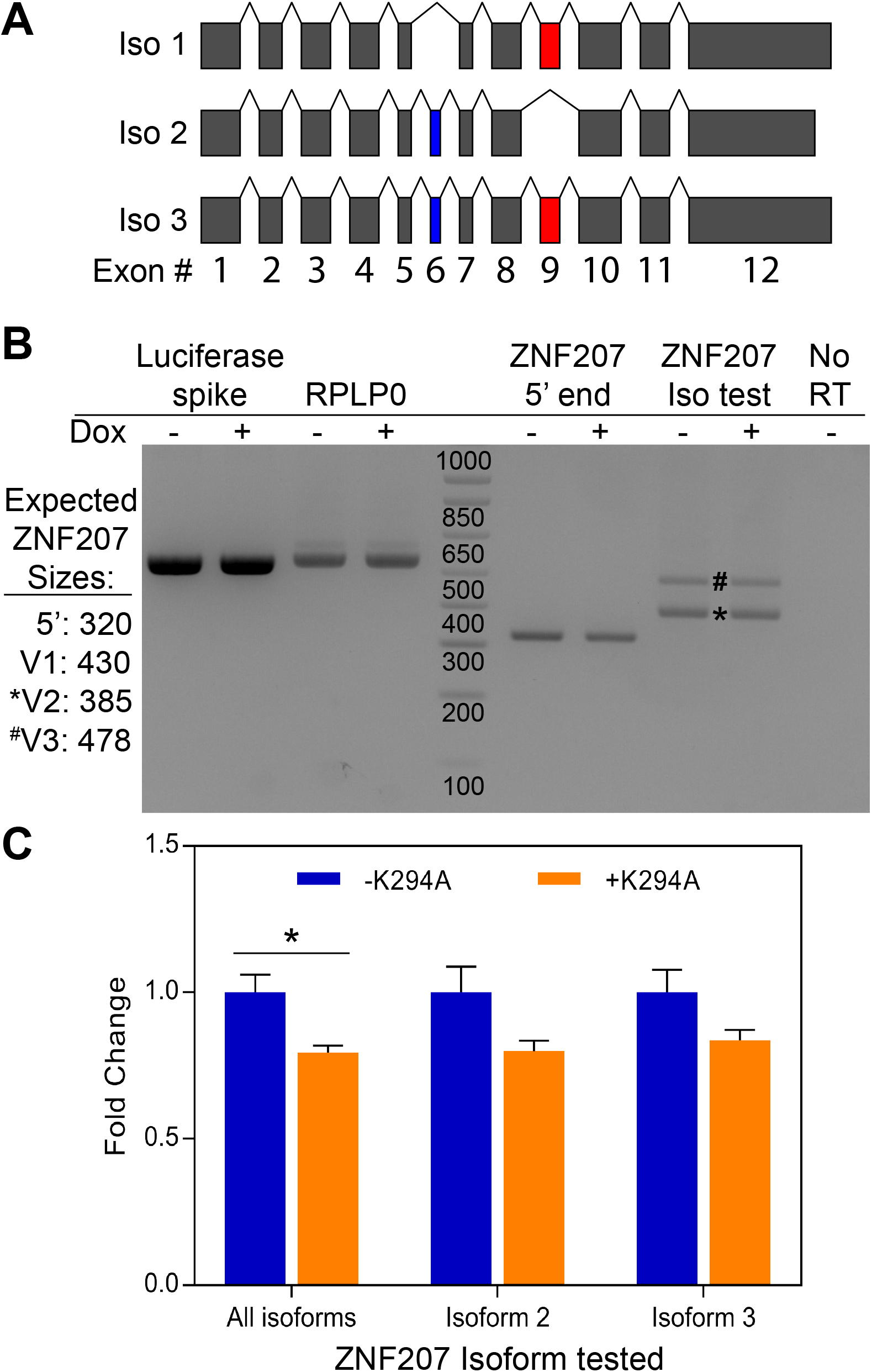
ZNF207 isoform 2 is a target of cytoplasmic mRNA recapping. (A) An exon map (exons drawn to scale) of the three most common ZNF207 splice isoforms. The isoform-distinguishing alternatively spliced exons, 6 and 9 are shaded (blue) and (red) respectively. **(B)** RNA was purified from cytoplasmic extracts of normal and K294-expressing cells, spiked with 0.1 ng of Luciferase RNA, reverse transcribed, and ZNF207 isoforms were assayed by standard RT-PCR. The predicted sizes of RT-PCR products for each isoform are shown to the left of the gel. Reactions using cDNA from a minus reverse transcriptase (No RT) sample (ZNF207 iso test PCR primers) and RT-PCR reactions targeting spiked luciferase and endogenous RPLP0 serve as controls. A representative agarose gel (three independent experiments) is shown. The positions of isoform 2-and 3-specific bands are marked by an * and # respectively. **(C)** cDNA remaining from above was assayed for expressed ZNF207 isoforms by qPCR (see methods). All data were normalized to spiked Luciferase mRNA and an endogenous control mRNA (Rplp0). The data shown are the means (± SEM) of three independent experiments each of which was analyzed in triplicate. The asterisk indicates a T-test p-value of <0.05.

Every read obtained in these 5’ RACE experiments targeting ITGB1 and SARS mRNAs spanned multiple exons; however, none of them could be conclusively mapped to an individual splice isoform. While the recovered clones didn’t map to a single splice isoform for either of these two mRNAs, nearly all (29/35) of the SARS clones mapped either to isoform 1 and 3, suggesting that the non-coding isoform 2 of SARS, NR_034072.1, is unlikely to be cytoplasmic capping target (Table S2).

### Identification of novel mRNA targets and uncapped 5’ ends for cytoplasmic mRNA recapping

Since uncapped ZNF207 mRNA was enriched in a single splice isoform, we reasoned that alternative splicing could help define which mRNAs were targeted by cytoplasmic capping. This hypothesis was tested by implementing a dual adaptor RACE (Figure 1C) strategy to identify novel 5’ RNA ends in an unbiased manner. As the guanylyl transferase activity of cytoplasmic capping enzyme is the penultimate step of cytoplasmic recapping, expressing K294A may result in a population of uncapped RNAs with 5’ diphosphate ends (Figure 1A) [5, 17]. Since 5’ RACE requires a 5’ monophosphate end, rSAP was used to remove any phosphate groups from the 5’ end of the RNA, and followed with a PNK treatment to return a single phosphate to any uncapped RNAs. It is important to note that the rSAP and PNK treatments have no effect on capped RNAs. In total, we cloned and sequenced 75 mRNA ends (34 in uninduced cells, and 41 in K294A expressing cells) mapping to 46 different mRNAs (Figure 4A, Table S3). As before, recovering uncapped mRNA ends in cells where cytoplasmic capping is unimpeded concurs with earlier results that showed stable uncapped mRNAs in normal cells [19]. Further, even with a limited number of clones, 6 mRNAs previously identified as cytoplasmic capping targets were recovered and validated (Figure 4A, Table S3) [19].

**Figure 4.**
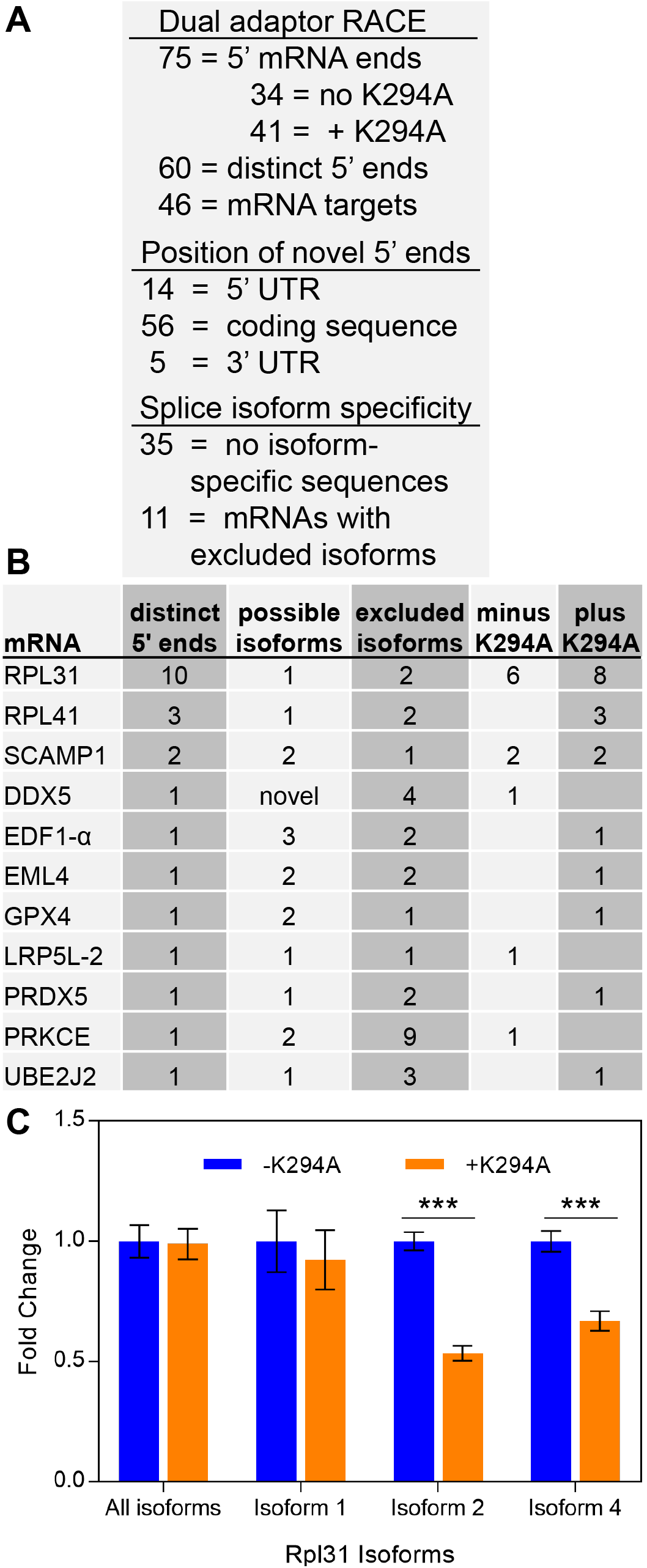
Dual adaptor RACE finds novel 5’ ends from truncated mRNAs. (A) A numerical summary of the novel 5’ mRNA ends discovered using the dual adaptor RACE approach. **(B)** A table listing the eleven mRNAs which included splice isoformspecific sequences. Also included are the number of 5’ ends observed, the number of possible and excluded isoforms and the whether the clones were observed in K294A-expressing (plus) or uninduced (minus) cells. **(C)** RNA was purified from cytoplasmic extracts of uninduced and K294-expressing cells, reverse transcribed, and Rpl31 isoforms were assayed by qPCR (see methods). All data were normalized to an external spike mRNA (Luciferase) and an endogenous control mRNA (Rplp0). The data shown are the means (± SEM) of three independent experiments each of which was analyzed in triplicate. *** indicates a T-test p-value of <0.0001.

A total of 60 new 5’ ends were identified, of which 39 were cloned only once (Figure 4A, Table S3). Of mRNAs identified multiple times, sequences mapping to RPL31 were the most common (14 clones, 6 in normal and 8 in +K294A cells), but ten distinct uncapped 5’ ends were cataloged for this mRNA (Table S3). Ten other mRNAs: DNAJB5 (5 clones, 1 end), NDUFA6 (3 clones, 1 end), RPL41 (3 clones, 1 end), DDX5 (3 clones, 3 ends), plus EEF1A1 (2 clones, 1 end), EMC6 (2 clones, 1 end), SCAMP1 (2 clones, 2 ends), SPARC (2 clones, 2 ends), TAF10 (2 clones, 1 end), and UBR1 (2 clones, 1 end) were cloned multiple times (Table S3).

### Most, but not all, recapping substrates could yield translatable ORFs when recapped

Most (59/75) of the novel 5’ ends reside within the 5’ UTR or the coding region of their host RNAs, suggesting that a truncated mRNA recapped at that position could be translated into an N-terminally truncated polypeptide. As 5’ UTR elements such as RNA secondary structures and upstream open reading frames are known to influence the translation of mRNAs, truncations in 5’ UTR sequences can also have large effects on the translational efficiencies of their mRNAs [24, 25]. Of the fourteen 5’ ends that mapped within 5’ UTRs, four mRNAs (eIF3D, HMCES, LRP5L-2, and LYSMD1) would have more than 150 bases truncated from the mRNA’s traditional 5’ UTR (Table S3). As the number of putative cytoplasmic recapping sites continues to expand, it will be interesting to see if different 5’ UTR sequence elements are preferentially retained by cytoplasmic recapping.

At the other extreme, five of these novel 5’ mRNA ends (DNAJB5, EEF1D, EMC6, NDUFA6, and RPL18A) were between 4 and 110 bases away from their stop codons, and four additional mRNAs (ATP6AP1, DAPK3, MTX3, and SPARC) had novel 5’ ends that were located entirely within the 3’UTR of the mRNA (Table S3). Since these nine truncated mRNAs begin close to-or after the stop codon, and do not contain obvious open reading frames, it is unlikely that they are translated into proteins or polypeptides. It is possible that such recapping targets could function as competing endogenous RNAs (ceRNAs). ceRNAs are becoming better characterized, and some have been shown to act as miRNA sponges to modulate the regulation of miRNA-targeted genes in certain contexts [26-29]. CD44, *VCAN*, and ESR1 mRNAs are just a few mRNAs whose 3’ UTRs have been shown to serve as ceRNAs that regulate the expression of other mRNAs [30-32].

### The truncated 5’ mRNA ends detected by dual adaptor RACE may be determined by alternative splicing

As above, nearly every cloned sequence spanned one or more splice junctions; although, in most cases, they could not differentiate between splice isoforms. However, in eleven cases (RPL31, RPL41, SCAMP1, DDX5, EDF1-α, EML4, GPX4, LRP5L-2, PRDX5, PRKCE, and UBE2J2) our clones contained splice junctions that excluded one or more isoforms (Figure 4B, Table S3). The result was most pronounced for RPL31 mRNA, where 11 of 14 newly cloned ends contained sequences that were restricted to isoform 1 (Table S3). As with the isoform-specific reads for ZNF207 mRNA above, the three ambiguous clones were too short to contain splice isoform defining sequences. Utilizing qPCR, we determined that the three best characterized protein coding Rpl31 isoforms were all expressed in U20S cells, although at substantially different levels (average efficiency-adjusted Cq values of 24.2, 30.3, and 27.5 for Isoforms 1, 2 and 4 respectively). The levels of Rpl31 mRNA isoforms were differentially affected by expressing K294A. While the total pool of Rpl31 mRNA and isoform 1 did not change significantly in response to K294A expression (Figure 4C), the two more poorly-expressed isoforms did decrease roughly two-fold.

Significant progress has been made in characterizing the proteins and mRNA targets of mammalian cytoplasmic mRNA recapping [33]. Nck1 nucleates the assembly of the cytoplasmic capping complex via interactions between its SH3 domains and capping enzyme [17]. RNMT is also part of the complex and methylates the newly capped transcripts making them identical to a nuclear-capped mRNAs [18]. A newly published paper has shown that RNA binding proteins can interact directly with capping enzyme and may recruit RNAs to the cytoplasmic capping complex [34]. As for the targets of cytoplasmic capping, a preliminary list of mRNAs has been identified [19], CAGE tags were shown to approximate the positions of uncapped 5’ mRNA ends [21], and uncapped cytoplasmic capping targets were shown to retain translatable poly(A) tails [20]. The work presented here reinforces and extends upon these earlier observations [19, 21]. Importantly, cytoplasmic capping also appears to have evolved independently in trypanosomes [35].

However, until now, there has been comparatively little progress in determining the rules that designated an mRNA for, or marked the position of, cytoplasmic mRNA recapping. The observation that only one splice isoform of ZNF207, and possibly other mRNAs, is targeted by recapping is a provocative one. It suggests that recapping may be determined by the sequences of the mRNAs themselves. This implies that cytoplasmic capping is a regulated process and that sequence elements that are included, excluded, or generated by alternative splicing may determine the occurrence and/or position of cytoplasmic recapping. While a possible link between alternative splicing and cytoplasmic capping is exciting, since only a small number of transcripts were cloned and each splice isoform-defining transcript appeared only a handful of times, a more in-depth analysis is still required. Importantly, this work also confirms that a population of mRNAs, the natively uncapped pool, exists in an uncapped, yet stable, form even when cytoplasmic capping is normal [19-21]. Collectively, these data offer the first insights into identifying an mRNA recapping signature sequence that not only selects mRNAs for recapping but may also mark the actual recapping sites.

## Acknowledgements

DLK conceived, designed, performed (or supervised) the experiments, and wrote the paper. MRB cloned, screened, and identified the novel 5’ ends for the dual adaptor RACE experiments and RA performed the RT-PCR and RT-qPCR experiments. MRB, RA, and DLK edited the manuscript. This work was supported by grant GM084177 from the National Institute of General Medical Sciences to Daniel R. Schoenberg and by startup funds provided by the Houston Methodist Research Institute (to DLK). The authors would like to thank Dr. Daniel Schoenberg his support of this work and his suggestions on the manuscript. MRB (undergraduate researcher) was supported by a Research Scholar award from The Ohio State University’s Office of Undergraduate Research and Creative Inquiry and by a Mayer’s Summer Research Scholarship from The Ohio State University’s College of Arts and Sciences. The OSU Genomics Shared Resource is supported by a grant from the National Cancer Institute (P30CA016058). The content is solely the responsibility of the authors and does not represent the official views of The Ohio State University, the Houston Methodist Research Institute, or the National Institutes of Health.

## Supplementary information

**Table S1: Oligonucleotides Used**

**Table S2: Targeted 5’ RACE experiments**

**Table S3: Dual adaptor RACE experiments**

**Figure S1. Most uncapped 5’-ends of truncated cytoplasmic capping-targeted ITGB1 and SARS mRNAs correspond to CAGE tags**. Schematics showing the positions of the novel 5’ ends cloned by 5’ RACE relative to full-length **(A)** ITGB1 and **(B)** SARS mRNAs. The green boxes in each schematic represent the coding sequence of the mRNA. The full-length transcript for each mRNA is indicated by m^7^G, and the nucleotide positions of the novel, 5’-truncated ends are shown to the left of each diagram. The number of times each 5’ end was cloned (blue box) and the number of CAGE tags mapping to that nucleotide position in the mRNA (grey box) are indicated to the right. The ^@^ denotes an exact 5’ RACE match to uncapped ends published in [21]Figure 1.

## List of Abbreviations

m^7^G: N7-methylguanosine cap
RNMT: RNA guanine-7 methyltransferase
NMD: nonsense-mediated decay
CAGE: capped analysis of gene expression
RACE: rapid amplification of cDNA ends
IPTG: Isopropyl β-D-1- thiogalactopyranoside
X-Gal: 5-bromo-4-chloro-3-indolyl-β-D-galactopyranoside
K294A: dominant negative cytoplasmic capping enzyme

